# Fluorescence Imaging-Based Discovery of Membrane Domain-Associated Proteins in *Mycobacterium smegmatis*

**DOI:** 10.1101/2021.02.17.431749

**Authors:** Corelle A. Z. Rokicki, James R. Brenner, Alexander Dills, Julius J. Judd, Jemila C. Kester, Julia Puffal, Joseph T. Wade, Todd A. Gray, Keith M. Derbyshire, Sarah M. Fortune, Yasu S. Morita

**Author notes:** Corelle A. Z. Rokicki, Covance, Madison, WI, USA. Julia Puffal, Department of Biochemistry and Molecular Biology, Rutgers University, NJ, USA. Julius Judd, Department of Molecular Biology and Genetics, Cornell University, Ithaca, NY, USA. Jemila C. Kester, Kaleido Biosciences, Lexington, MA, USA. To whom correspondence should be addressed: Department of Microbiology, University of Massachusetts, 639 North Pleasant Street, Amherst MA 01003, USA.

## Abstract

Mycobacteria spatially organize their plasma membrane, and many enzymes involved in envelope biosynthesis associate with a membrane compartment termed the intracellular membrane domain (IMD). The IMD is concentrated in the polar regions of growing cells and becomes less polarized under non-growing conditions. Because mycobacteria elongate from the poles, the observed polar localization of the IMD during growth likely supports the localized envelope biosynthesis. While we have identified more than 300 IMD-associated proteins by proteomic analyses, only a handful of these have been verified by other experimental methods. Furthermore, we speculate that some IMD-associated proteins may have escaped proteomic identification and remain to be identified. Here, we visually screened an arrayed library of 523 *Mycobacterium smegmatis* strains each expressing a Dendra2-FLAG-tagged recombinant protein. We identified 29 fusion proteins that showed fluorescence patterns similar to those of IMD proteins and, consistent with this co-localization, we had previously identified 20 of these using a proteomics approach. Of the nine remaining IMD candidate proteins, three were confirmed to be associated with the IMD while some others appear to be lipid droplet-associated. Taken together, our newly devised strategy is effective in verifying the IMD association of proteins found by proteomic analyses, while facilitating the discovery of additional IMD-associated proteins.

**Importance:** The intracellular membrane domain (IMD) is a membrane subcompartment found in *Mycobacterium smegmatis* cells. Proteomic analysis of purified IMD identified more than 300 proteins, including enzymes involved in cell envelope biosynthesis, that likely contribute to the function of the IMD. How can we find more IMD-associated proteins that escaped proteomic detection? Here, as an alternative approach, fluorescence microscope images of 523 proteins were screened to identify IMD-associated proteins. We confirmed the IMD association of previously identified proteins and discovered three additional proteins associated with the IMD. Together, subcellular fractionation, proteomics, and fluorescence microscopy form a robust combination to more rigorously define IMD proteins, which will aid future investigations to decipher the synthesis, maintenance and functions of this membrane domain.

## Introduction

The cell envelope of mycobacteria comprises the plasma membrane, cell wall and outer membrane. The diderm structure is unusually complex with a peptidoglycan layer covalently linked to an arabinogalactan layer, which is further modified by mycolic acids. Mycolic acids form a part of the outer membrane, which contributes to creating a characteristic waxy outer coating (1–5).

Mycobacteria must retain this complex cell envelope as they grow and extend their rod-shaped cell. New components for cell envelope extension are added at the subpolar regions (6–12). The intracellular membrane domain (IMD) is distinct from the conventional plasma membrane, and is found enriched in the subpolar regions of actively growing cells (13, 14). The IMD is spatially positioned to support the polar cell wall synthesis and growth, and its subpolar enrichment is reduced when the cells stop growing (15, 16). The IMD can be biochemically isolated from the cell lysate of *Mycobacterium smegmatis* by sucrose density gradient sedimentation (14). During fractionation, the IMD sediments to a region of lighter density than the plasma membrane. The plasma membrane is purified in tight association with the cell wall components, and the complex is designated as plasma membrane-cell wall (PM-CW). Proteomic analysis of the IMD revealed that more than 300 proteins potentially associate with the IMD. Well-characterized examples of the IMD proteins include cell envelope biosynthetic enzymes such as PimB’, Ppm1, and GlfT2 (13). PimB’ is an essential mannosyltransferase that catalyzes the transfer of a second mannose to form phosphatidylinositol dimannosides, a component of the cell envelope (17, 18). Ppm1 is a subunit of the polyprenol phosphate mannnose (PPM) synthase involved in synthesizing lipid-linked mannose donors, and is another essential enzyme for the synthesis of mannose-containing glycolipids (19–22). GlfT2 is a galactosyltransferase that extends the galactan chain of the arabinogalactan layer (23, 24). The identification of these biosynthetic enzymes, among others involved in cell envelope biosynthesis, highlights the critical roles of the IMD in cell envelope biosynthesis.

Although over 300 putative IMD associating proteins have been identified, we anticipate that there will be additional IMD proteins that were not identified by proteomics due to its inherent limitations such as inefficient detection of membrane proteins. In this study, we took advantage of a recently established Mycobacterial Systems Resource. This resource includes a library of *Mycobacterium smegmatis* strains, each expressing one of 1118 genes in the core genome of five *Mycobacterium* species as a recombinant protein tagged with C-terminal Dendra2-FLAG tag. We visually screened Dendra2 fluorescence patterns of 523 images for IMD-like fluorescence and identified 29 putative IMD proteins. Among them, 20 were previously found in the IMD proteome; we provide biochemical validation of 5 previously found IMD proteins as well as three new IMD proteins.

## Results

### Screening of fluorescence microscope images identifies candidate IMD-associated proteins

As an alternative approach to discover additional IMD-associated proteins, we screened fluorescence microscopy images of 523 *M. smegmatis* strains each expressing a fusion protein, which is tagged with Dendra2-FLAG at the C-terminus (**Table S1**) (see *Materials and Methods*). We defined the IMD-like fluorescence pattern as fluorescence enrichment in the polar regions with weaker fluorescence patches along the sidewalls (13). Through visual screening, we identified 29 potential strains, showing fluorescence patterns of Dendra2-FLAG-tagged proteins that are similar to the typical localization patterns of IMD-associated proteins (**Fig. S1**). The identified proteins were then cross-referenced against the IMD proteome. We found that 20 of the 29 proteins were those previously found by the proteomic analysis (**Table 1**). One of the remaining 9 proteins, MenA (MSMEG_0988), was not listed in our proteome, but we have shown in a separate study that it is a PM-CW-associated protein (25). Therefore, we considered this a false-positive, and did not investigate this protein further. Among the 8 new candidates, five proteins were previously found in the PM-CW proteome (13), whereas the remaining three were not found in either IMD or PM-CW proteome (**Table 1**).

**Table 1.** IMD protein candidates identified from visual screening of fluorescence microscope images.

| MSMEG | Rv | Gene | Annotation | Proteome | Biochemical (Ref) |
| --- | --- | --- | --- | --- | --- |
| 0825 | Rv0422c | thiD | phosphomethylpyrimidine kinase | PM-CW | IMD (this study) |
| 0861 <sup>*1</sup> | Rv0437c | psd | phosphatidylserine decarboxylase | IMD | IMD (14) |
| 0863 | Rv0439c |  | short chain dehydrogenase | IMD |  |
| 0876 | Rv0068 |  | short chain dehydrogenase | IMD | IMD (this study) |
| 0949 | Rv0505c |  | HAD-superfamily hydrolase | IMD | C <sup>*2</sup> (this study) |
| 0972 | Rv0527 | ccdA | cytochrome C biogenesis protein transmembrane region | NF <sup>*3</sup> | IMD (this study) |
| 0988 <sup>*4</sup> | Rv0534c | menA | 1,4minusdihydroxyminus2minusnaphthoate octaprenyltransferase | NF | PM-CW (25) |
| 1011 | N/A |  | short chain dehydrogenase | IMD |  |
| 1055 | N/A |  | hexapeptide transferase | IMD |  |
| 1115 | Rv0558 | menG | ubiquinone/menaquinone biosynthesis methyltransferase | IMD | IMD (25) |
| 1352 | Rv0646c |  | alpha/beta hydrolase | IMD |  |
| 1954 | Rv3197 |  | ABC transporter | PM-CW | C (this study) |
| 2080 | Rv3140 | fadE23 | Acyl-CoA dehydrogenase | IMD |  |
| 2329 <sup>*5</sup> | Rv3038c |  | UbiE/COQ5 family methyltransferase | IMD | C (LD) (this study) |
| 2335 <sup>*5</sup> | Rv3034c |  | hexapeptide transferase | IMD | IMD (this study) |
| 2698 | Rv2740 |  | hypothetical protein MSMEG_2698 | IMD |  |
| 2934 | Rv2611c | patA | lipid A biosynthesis lauroyl acyltransferase | IMD | IMD (this study) |
| 2975 | Rv2581c |  | Metallo-beta-lactamase | NF |  |
| 3863 | Rv2061c |  | pyridoxamine 5'-phosphate oxidase | IMD |  |
| 4198 | Rv2139 | pyrD | dihydroorotate dehydrogenase 2 | IMD | IMD (this study) (13) |
| 4227 | Rv2153c | murG | UDP diphospho-muramoyl pentapeptide beta-N-acetylglucosaminyltransferase | IMD |  |
| 4254 | Rv2187 | fadD15 | AMP-binding protein | PM-CW | C (this study) |
| 4284 | Rv2216 |  | hypothetical protein MSMEG_4284 | IMD |  |
| 4479 | Rv2342 |  | hypothetical protein MSMEG_4479 | NF | IMD (this study) |
| 4632 | Rv2449c |  | saccharopine dehydrogenase | IMD |  |
| 4722 | Rv2509 | cmrA | Short-chain dehydrogenase | PM-CW | C (LD) (this study) |
| 6385 | Rv3791 |  | Short chain dehydrogenase | IMD |  |
| 6403 | Rv3808c | glfT2 | bifunctional udp-galactofuranosyl transferase glfT | IMD | IMD (this study) (13) |
| 6928 | Rv3909 |  | hypothetical protein MSMEG_6928 | PM-CW | PM-CW (this study) |

<sup>\*1</sup> Gray shade indicates the proteins that were previously found in the IMD proteome.
<sup>\*2</sup> C, cytoplasm. This fraction may include lipid droplets (LD). See text for details.
\*3 NF, not found.
\*4 This protein has been previously characterized to be a PM-CW protein [Puffal 2018], and was not investigated further in this study.
\*5 MSMEG\_2329 and 2335 are identical to MSMEG\_1049 and 1055 at the amino acid level, respectively, and are both found in the IMD proteome.

### Confirmation of previously identified IMD proteins

Among the 20 previously identified IMD proteins, GlfT2, PyrD, MenG and MurG have already been examined by biochemical and fluorescence image analysis (8, 13, 14, 25), establishing the characteristic features of IMD-associated proteins. We chose two proteins, GlfT2 (MSMEG_6403) and PyrD (MSMEG_4198), to confirm that the Dendra2-FLAG tag fusions are biochemically associated with the IMD fraction. GlfT2, a galactosyltransferase involved in galactan synthesis, has been previously studied by three different fusion proteins: 1) GlfT2-HA expressed from a strong heterologous promoter; 2) mTurquoise-GlfT2-FLAG expressed from a strong heterologous promoter; and 3) HA-mCherry-GlfT2 expressed from its native locus (13). In addition, GlfT2-GFP is enriched in the subpolar region of the cell, consistent with its IMD classification (8).

The GlfT2-Dendra2-FLAG, described in this report, represents the fifth fusion construct to be tested for its subcellular localization. To examine the subcellular localization, we grew the strain expressing GlfT2-Dendra2-FLAG to log phase, lysed the cells by nitrogen cavitation, and subjected the lysate to sucrose density gradient fractionation. Equal volumes of each fraction were separated by SDS-PAGE and the fusion protein was detected by western blotting. We confirmed that GlfT2-Dendra2-FLAG, like other previously tested fusion constructs, was at the predicted molecular weight of 100.2 kDa, and specifically associated with the IMD (**Fig. 1A**). We defined the IMD enrichment as follows: 1) at least 60% of the total band intensity across the entire gradient fractions is enriched in the IMD fractions (Fr. 3-6); and 2) the protein is enriched more than two-fold in the IMD fractions (Fr. 3-6), relative to the cytoplasmic fractions (Fr. 1-2) and the PM-CW fractions (Fr. 8-11). Quantification of the western blot bands indicated that 99.6% of GlfT2-Dendra2-FLAG was associated with the IMD (**Fig. 1C**), establishing this fusion protein as a clear example of an IMD-associated protein.

**Figure 1.**
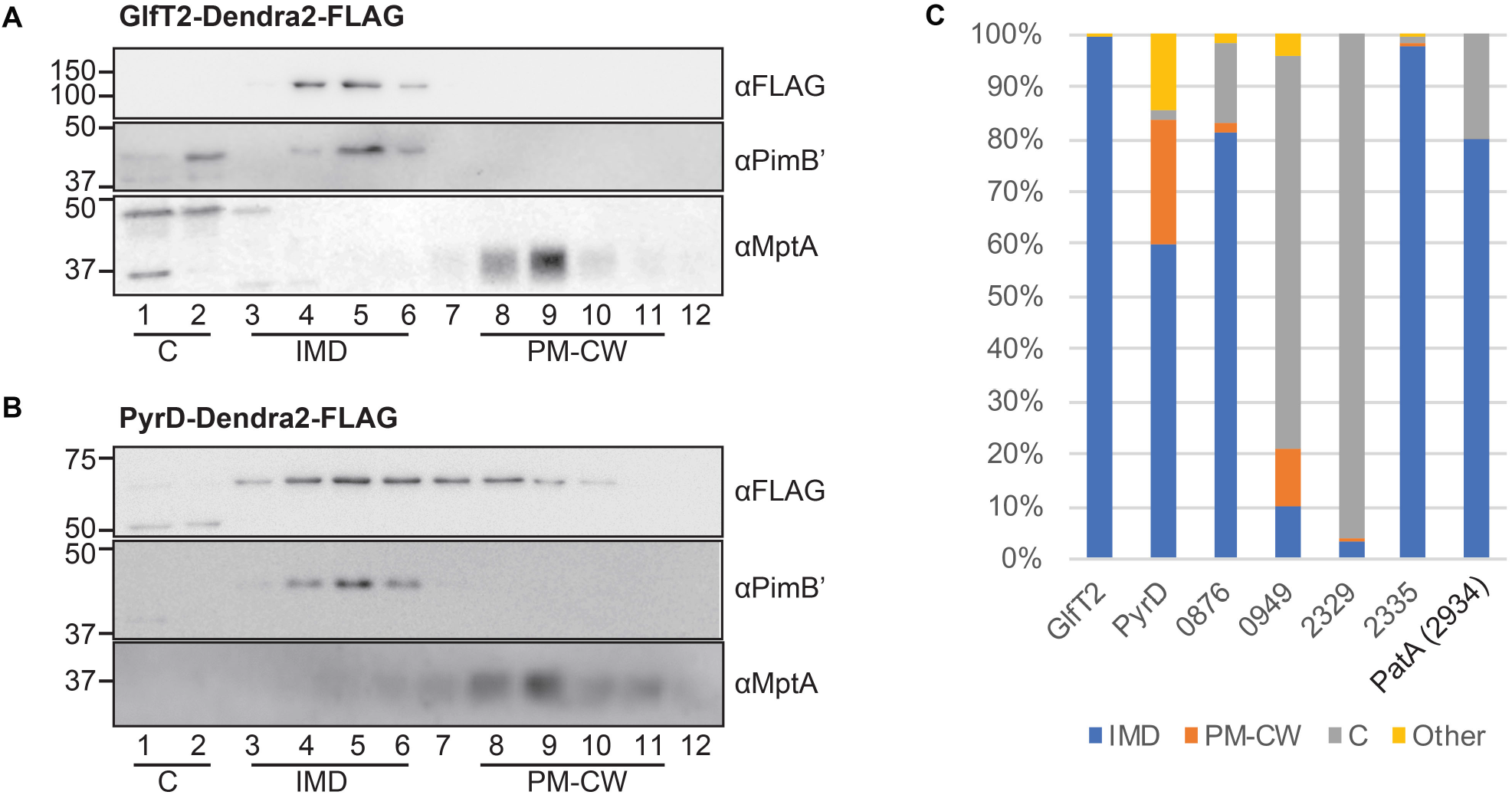
Validation of known IMD protein candidates by density gradient fractionation. Among IMD protein candidates identified by the image screening, seven proteins, which were previously found in the IMD proteome, were examined by sucrose density gradient fractionation. **(A)** Density gradient fractionation of GlfT2-Dendra2-FLAG. An equal volume of each gradient fraction was separated by SDS-PAGE, and the FLAG epitope was detected by anti-FLAG western blotting. PimB’ (41.1kDa), an IMD marker; MptA (54.3 kDa), a PM-CW marker. Apparent molecular weight of MptA on SDS-PAGE gel is about 40 kDa (32, 34). **(B)** Density gradient fractionation of PyrD-Dendra2-FLAG. **(C)** Quantification of western blot bands shown in Fig. 1A, 1B, and S2. Fr. 1-2, cytoplasm (C); Fr. 3-6, IMD; Fr. 8-11, PM-CW; Fr. 7 and 12, other.

The IMD association of PyrD, a dihydroorotate reductase involved in pyrimidine synthesis, has also been biochemically verified by the specific association of PyrD-HA with the IMD fractions of a sucrose gradient (13). Consistent with the previous finding, PyrD-Dendra2-FLAG was detected in the IMD fraction (**Fig. 1B)**. Quantification of PyrD-Dendra2-FLAG bands indicated that 60.0% was found in the IMD, and this enrichment was greater than 2-fold in comparison to the cytoplasm (1.75%) and the PM-CW (23.9%) (**Fig. 1C**). Therefore, we conclude that PyrD is an IMD-associated protein. Previously, we showed the IMD enrichment of PyrD-HA to be 78% (13), indicating that the addition of Dendra2-FLAG, a relatively large fluorescent protein tag, may have interfered with efficient localization of this protein to the IMD.

In addition to these two previously characterized proteins, we examined the biochemical partitioning of 5 additional proteins, MSMEG_0876 (short chain dehydrogenase), MSMEG_0949 (HAD-superfamily hydrolase), MSMEG_2329 (UbiE/COQ5 family methyltransferase), MSMEG_2335 (hexapeptide transferase), and MSMEG_2934 (PatA), that were previously found in the IMD proteome, but were not investigated further. The precise functions of these proteins are unknown except PatA, which is an acyltransferase involved in the synthesis of phosphatidylinositol mannosides (26). Our previous study suggested that PatA is likely an IMD-associated enzyme because a cell-free enzymatic assay showed the enriched production of AcPIM2 in the IMD fraction (14). However, there has been no biochemical confirmation of the protein enrichment in the IMD. As shown in **Fig. 1C and S2**, PatA and two other proteins, MSMEG_0876 and MSMEG_2335, were highly enriched in the IMD fractions of the density gradient. In contrast, two proteins, MSMEG_0949 and MSMEG_2329, were found to be more enriched in the cytoplasmic fraction (Fr. 1-2) (**Fig. 1C and S2**) (see below and Discussion). Taken together, microscopic and biochemical analyses of Dendra2-FLAG fusion proteins are effective in evaluating the IMD associations of proteins that are previously predicted to be IMD-associated.

### Verifying the expression of newly identified IMD candidates

Next, we examined the eight proteins that were not identified as an IMD protein in our previous proteomic analysis. We grew these 8 strains, prepared cell lysates, and analyzed the molecular weights of the fusion proteins by SDS-PAGE and western blotting. The western blot image of the 8 proteins demonstrated that MSMEG_0972 (CcdA), MSMEG_4254 (FadD15), MSMEG_4479, and MSMEG_6928 were at or near their expected molecular weights (**Fig. 2A**). The apparent molecular weights of MSMEG_0825 (ThiD), MSMEG_1954, and MSMEG_4722 (CmrA) were smaller than those expected. In the cases of MSMEG_1954 and CmrA, there is a fainter band that migrated near the expected molecular weight, suggesting that the more intense band could be a stable degradation product. In addition, we could not detect MSMEG_2975 by this method. As an alternative approach, we detected Dendra2 protein by fluorescence imaging of non-denaturing semi-native-PAGE. As shown in **Fig. 2B**, all of the proteins showed similar migration patterns to those observed by standard SDS-PAGE and western blot (compare with **Fig. 2A**), except that ThiD and MSMEG_4479 migrated slightly faster (likely proteolytic products of the Dendra2 protein). Interestingly, we could detect a faint MSMEG_2975-Dendra2-FLAG band on the semi-native gel, even though it was not detected by western blotting. In conclusion, these results indicate that all of the strains express the target protein with a functional Dendra2 tag and the apparent molecular weight on SDS-PAGE is mostly consistent with the expected size of the full-length fusion protein.

**Figure 2.**
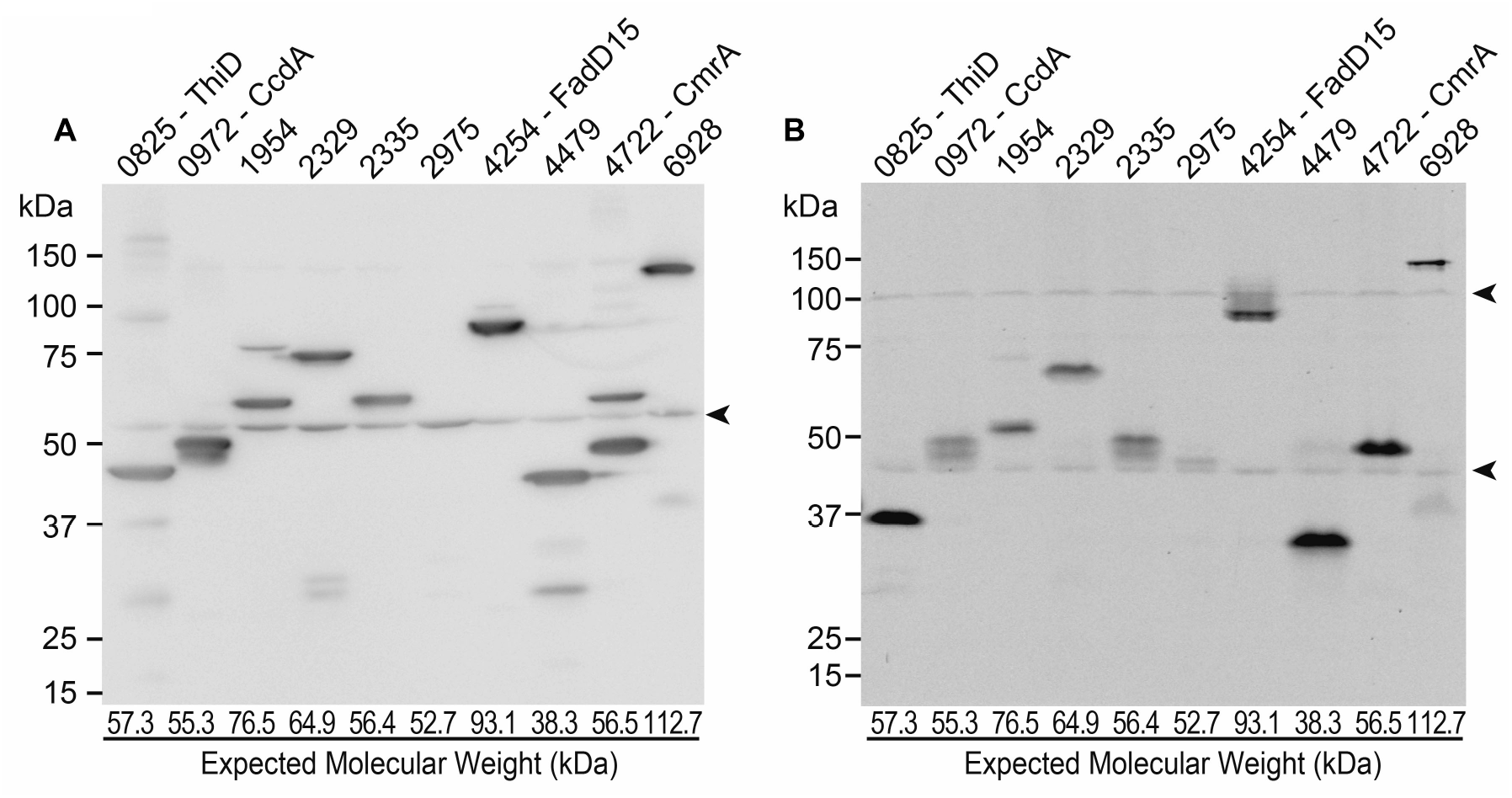
Western blot analysis of newly discovered IMD protein candidates. **(A)** Crude cell lysates of the 8 new IMD protein candidates (as well as MSMEG_2329 and MSMEG2335) were analyzed by SDS-PAGE, and FLAG epitope was detected by western blotting. Expected molecular weight of each protein was indicated at the bottom of the lane. **(B)** Crude cell lysates were analyzed by PAGE under a non-denaturing condition and Dendra2 fluorescence was detected in gel by a fluorescence imager. Arrowheads, non-specific proteins.

### MSMEG 4479 is an IMD-associated protein

Among the 8 candidates that were not found in the IMD proteome, we realized that MSMEG_4479 was missing from the list of annotated genes in the genome sequence (NC_018289, MSMEI_ locus tag prefix), which was used in our proteomic analysis (13). There are two frequently used complete genome sequences for *M. smegmatis* mc^2^155 strains in the NCBI Reference Sequence Database: NC_008596 submitted by TIGR on 11/20/2006 and NC_018289 submitted by the Institute of Genetics and Molecular and Cellular Biology (IGMCB) on 8/3/2012. There are 6716 and 6693 protein sequences annotated in the TIGR and IGMCB sequences, respectively. MSMEG_ is the locus tag prefix for the TIGR sequence, and when we cross-referenced MSMEG_4479 against the IGMCB sequence, we could not find the corresponding gene with the MSMEI_ locus tag prefix. We retrieved the genome region from these two complete sequences and searched for open reading frames. The DNA sequence was completely identical between the two sequences in the region 4000 bp upstream and downstream of MSMEG_4479 gene (**Fig. S3A**), however, the annotation of MSMEG_4479 is missing in the IGMCB sequence.

We reanalyzed our previous proteomic data using the TIGR sequence (NC_008596) as a reference genome. We found a normalized iBAQ value of 6.43×10^8^ for MSMEG_4479 from the immunoprecipitated IMD vesicles, which was 700 times more enriched than the non-specific binding in a negative control (**Fig. S3B**). The iBAQ value and the IMD enrichment were well above our previous threshold (13). Thus, our proteomic analysis is in agreement with the co-localization pattern suggesting that MSMEG_4479 is an IMD protein.

Next, we examined the subcellular localization of MSMEG_4479-Dendra2-FLAG by density gradient fractionation. The fusion protein was separated by SDS-PAGE and detected by western blotting (**Fig. S4**). We measured the band intensity of the fusion protein from each fraction and determined the relative distribution across the density gradient (**Fig. 3**). The enrichment of MSMEG_4479 in the IMD was 67.4%, and the protein enrichment in the cytoplasm and the PM-CW was less than 10%. Therefore, we biochemically confirmed MSMEG_4479 as an IMD protein.

**Figure 3.**
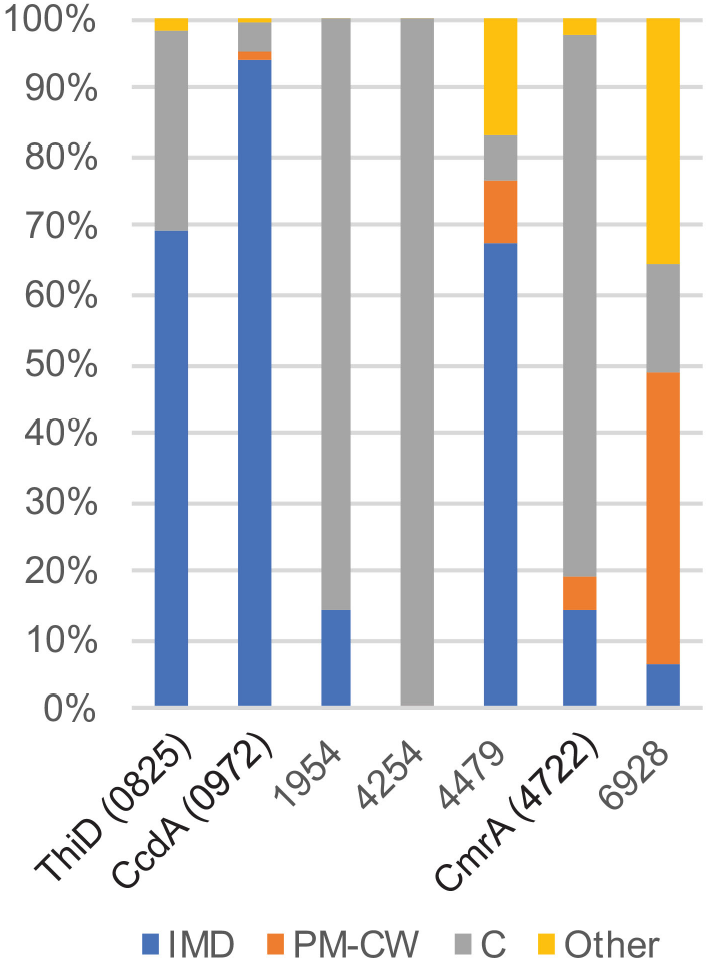
Subcellular fractionation of newly discovered IMD protein candidates. Western blot bands of each protein, as shown in Fig. S4, were quantified across density gradient fractions as described in Fig. 1C. ThiD, CcdA and MSMEG_4479 were confirmed as IMD proteins (see text for details).

### The IMD association of other candidate proteins determined by density gradient

We examined the remaining six new candidates by sucrose density gradient (**Fig. 3 and S4**). Note that MSMEG_2975 was not further analyzed because it was poorly expressed and co-migrated with a non-specific fluorescent protein on the gel (see **Fig. 2**). Based on the definition of IMD association described above, we classify ThiD and CcdA as two additional IMD-associated proteins.

ThiD is a phosphomethylpyrimidine kinase involved in thiamine biosynthesis (27). As mentioned above, the apparent molecular weight of ThiD was smaller than the expected molecular weights (see **Fig. 2**). The cell lysates were prepared by a bead-beating cell disruptor for this initial experiment. Vortical flow of beads generates a shear force that efficiently lyses the cell, but also produces heat due to the friction of beads. In contrast, for sucrose gradient fractionation, the cell lysates were prepared by nitrogen cavitation, which does not generate heat. Interestingly, using nitrogen cavitation lysate, ThiD migrated with an apparent molecular weight that is much closer to its predicted size of 57.3 kDa (**Fig. S4**), suggesting that the full-length protein might be heat sensitive. The fractionation pattern of ThiD indicates that it is enriched in the IMD (69.6%) relative to the PM-CW (0%) and the cytoplasm (28.8%) (**Fig. 3**). Therefore, we concluded that this protein is an IMD-associated protein.

CcdA is a putative reductase that is predicted to regenerate the reduced form of the periplasmic thioredoxin CcsX, which is essential for the maturation of cytochrome C (28). The amino acid sequence of CcdA suggests that it is a polytopic transmembrane protein. When analyzed by density gradient sedimentation, CcdA-Dendra2-FLAG was highly enriched in the IMD (94.4%) (**Fig. 3**). To further confirm the IMD localization of CcdA, we expressed it in a strain, in which the endogenous gene encoding GlfT2 is replaced with a fusion encoding HA-mCherry-GlfT2 (13). We found that CcdA co-localized with HA-mCherry-GlfT2, a well-established IMD marker (**Fig. 4**).

**Figure 4.**
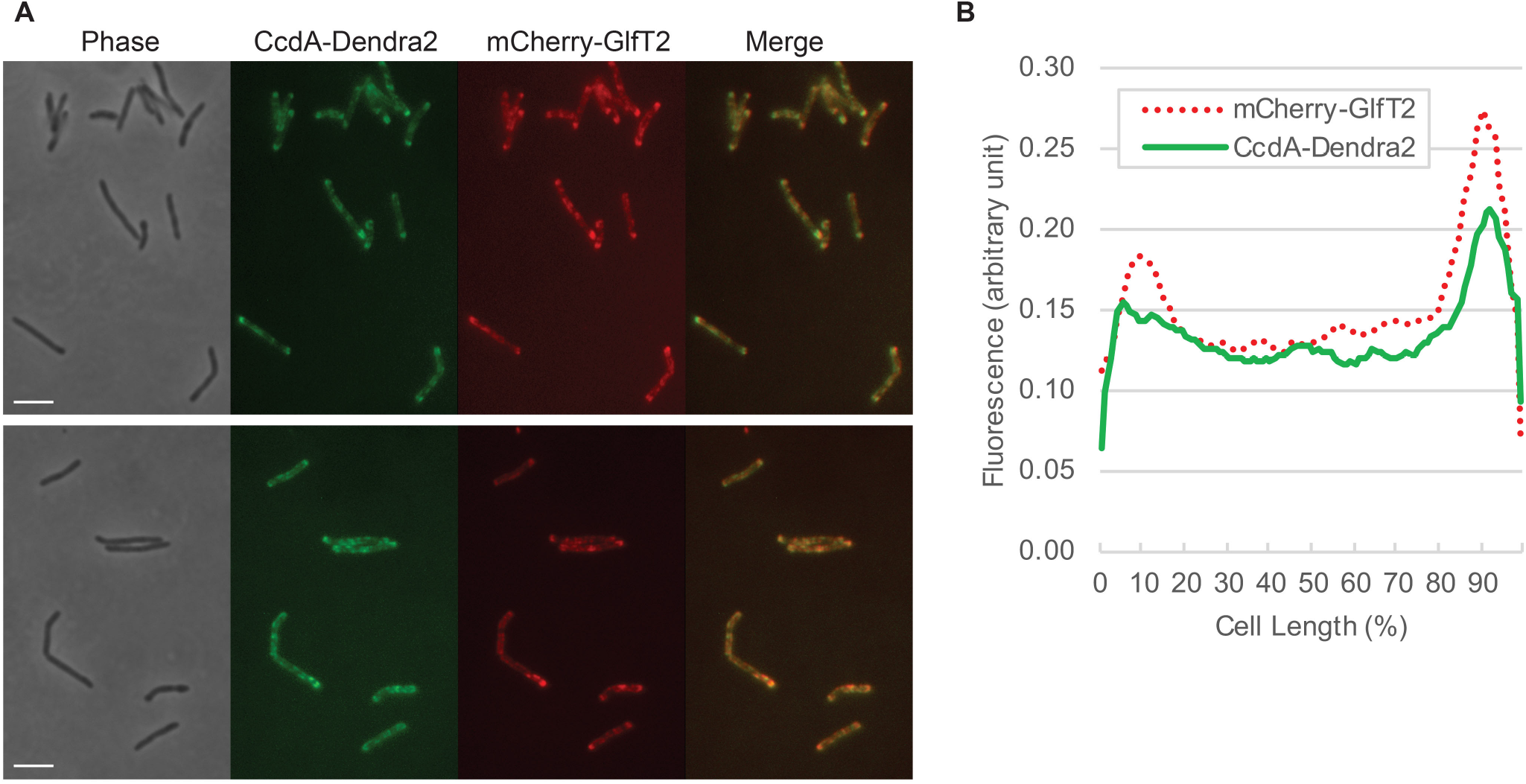
Co-localization of CcdA-Dendra2-FLAG and HA-mCherry-GlfT2 visualized by fluorescence microscopy. **(A)** HA-mCherry-GlfT2 is expressed from the endogenous locus and previously confirmed as an IMD marker (13). Two different fields are shown as representative images. Bar = 4 μm. **(B)** Fluorescence intensity profile along the long axis of the cell. Cell length is shown in percent and cells were aligned so that a more intense pole is positioned to the right (100%). Average intensity profiles of 14 cells are shown.

The other four proteins, MSMEG_1954, FadD15, CmrA, and MSMEG_6928, were previously assigned to the PM-CW by proteomic analysis (13), However, our analysis shows that MSMEG_6928 is the only protein that is partially (~40%) associated with the PM-CW in the density gradient fractionation (**Fig. 3 and S4**). The other three proteins were found in the cytoplasmic fraction. Similar to ThiD, full-length CmrA protein was predominant after lysis by nitrogen cavitation (**Fig. S4**). CmrA did not associate with the IMD at all, and was highly enriched in the cytoplasmic fraction (**Fig. 3 and S4**).

However, a small amount of the protein is detectable in the PM-CW fractions, suggesting that this protein may partially associate with the PM-CW. In contrast to ThiD and CmrA, both full length (76.5 kDa) and truncated (~50 kDa) versions of MSMEG_1954 protein remained detectable even after nitrogen cavitation cell lysis and both forms of MSMEG_1954 were enriched in the cytoplasmic fraction (**Fig. S4**). Therefore, biochemical characterizations were not consistent with the prediction that these proteins are IMD-associated based on fluorescence imaging (see Discussion).

### MSMEG_2329 and CmrA partially associate with lipid droplets

It is striking that several proteins that were previously (and here by co-localization) predicted to be associated with either the IMD or the PM-CW stayed at or near the top of the gradient (fractions 1-2), where cytoplasmic proteins are normally enriched (13). The five proteins we showed here to be localized in the cytoplasm (MSMEG_0949, 1954, 2329, FadD15, and CmrA), were previously identified in a lipid droplet proteome (29). We chose two of these proteins, MSMEG_2329, and CmrA for further investigation. Lipid droplets are buoyant, and density gradient flotation is a well-established method to identify lipid droplet proteins (29, 30). Therefore, we placed the lysate at the bottom of a 20-50% sucrose gradient and fractionated by isopycnic ultracentrifugation to examine if proteins float to the top of the gradient. As shown in **Fig. 5**, a small fraction (14 and 3%, respectively) of MSMEG_2329 and CmrA floated to the top of the gradient, although the majority of the proteins stayed at or near the bottom of the gradient. In the case of CmrA, a significant fraction (~67%) of the protein floated and was enriched in fractions 8-12, which corresponded to PM-CW (as defined by the marker MptC; see Discussion). Taken together, our data suggest that MSMEG_2329 and CmrA are at least partially associated with lipid droplets as previously predicted by lipid droplet proteome analysis.

**Figure 5.**
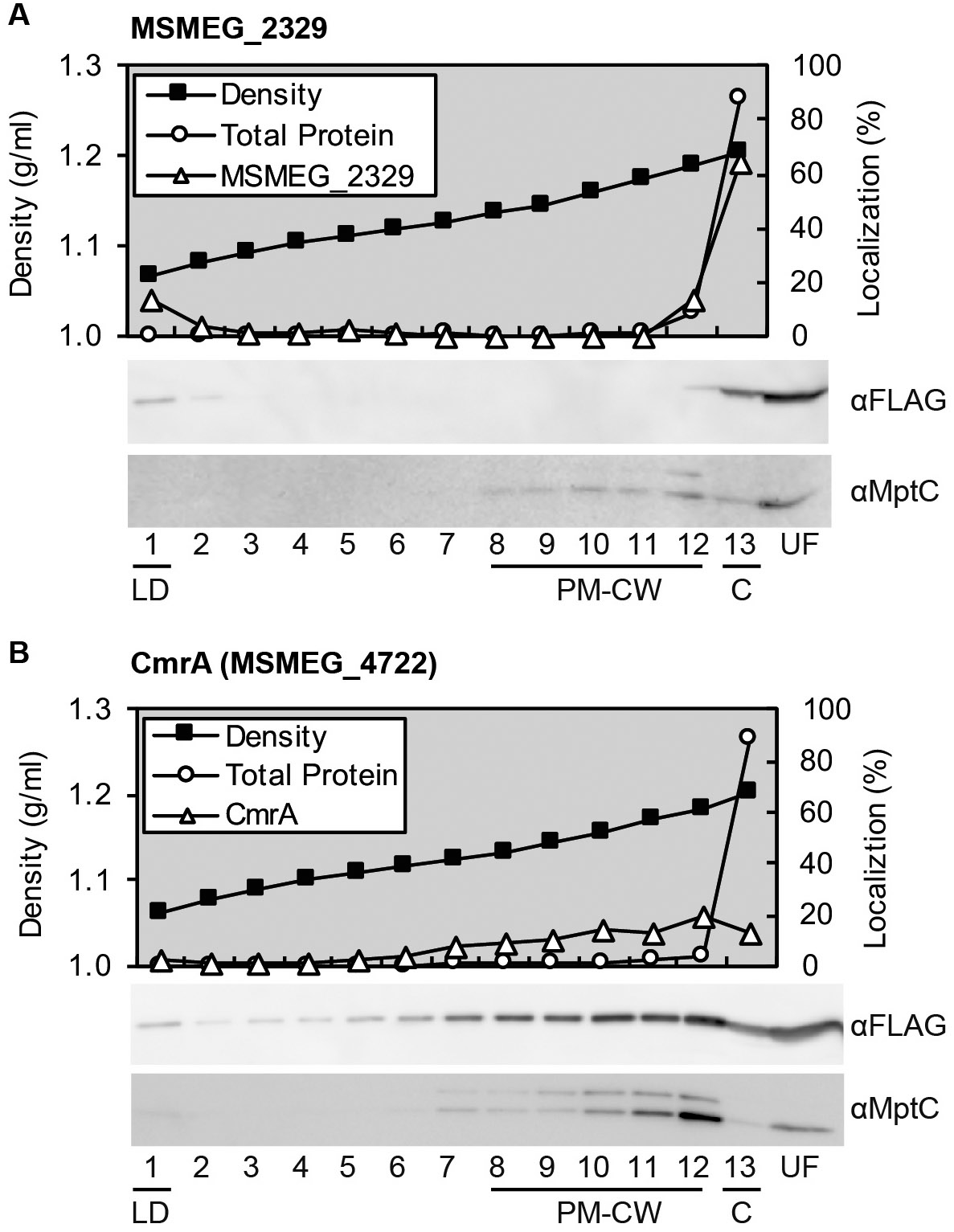
Density gradient flotation analysis of MSMEG_2329 and CmrA. **(A)** Lysate of *M. smegmatis* cells expressing MSMEG_2329 placed to the bottom of sucrose gradient and fractionated by isopycnic ultracentrifugation. Relative localization of total proteins was determined by a BCA assay, while that of MSMEG_2329 was calculated from density measurement of western blot bands using Fiji (35). Total proteins essentially represent bulk cytoplasmic proteins that remain at the bottom fraction (Fr. 13). Density was determined by refractive index. For western blotting, equal volume of each fraction was separated by SDS-PAGE, and protein bands were identified by anti-FLAG (αFLAG) or anti-MptC (αMptC) antibody. MptC is a PM-CW marker, which floated to fractions 8-12. A portion of MSMEG_2329 floated to the top fraction, which correspond to the lipid droplet fraction (Fr. 1). **(B)** Lysate of cells expressing CmrA was fractionated and analyzed as described in panel A.

## Discussion

In our previous study, we purified the IMD by density gradient and immunoprecipitation, and analyzed it by proteomics. We reported 309 proteins as IMD-associated proteins. In the current study, we used the Mycobacterial Systems Resource to visually screen nearly 600 images of fluorescent protein localizations in *M. smegmatis* and identified 29 IMD candidates. Among them, 20 proteins were previously reported as a part of the IMD proteome, while 3 proteins were newly validated as IMD-associated proteins. Thus, fluorescence image screening is an independent method to identify additional IMD-associated proteins that escaped proteomic discovery.

We analyzed 7 out of 20 previously identified IMD proteins to validate the subcellular localization biochemically. Five of them were not characterized previously, and among them MSMEG_0876, MSMEG_2335, and PatA were enriched in the IMD fraction after density gradient fractionation. In contrast, two proteins, MSMEG_0949 and MSMEG_2329, appeared cytoplasmic and were not associated with the IMD. It remains possible that these proteins are associated with the IMD *in vivo*, but were released from the membrane upon cell disruption. Alternatively, they may be cytoplasmic or lipid-droplet-associated (see below). Determining the true subcellular localization of these proteins requires more in-depth studies of individual proteins. Taken together, the image screening is effective in identifying proteins that could potentially be a false positive of proteomic analysis, highlighting the importance of combining microscopic and biochemical validations for proteomics data.

Strikingly, we could identify only 3 additional IMD proteins (ThiD, CcdA and MSMEG_4479) among 9 new candidates. Identification of MSMEG_4479 highlights the importance of accurate gene annotation in the database we use for proteomic analysis. ThiD and CcdA are the only two truly new IMD proteins that were not identified by the proteomic analysis. CcdA is an integral membrane protein with 6 predicted transmembrane segments. We suspect that the hydrophobic nature of the protein may be one of the reasons why this protein has escaped detection by proteomics.

We found five proteins to be cytoplasmic: FadD15, and CmrA, MSMEG_0949, MSMEG_1954, and MSMEG_2329. Interestingly, all of these proteins were previously associated with a lipid droplet proteome (29). We tested CmrA and MSMEG_2329 using density gradient flotation and both partially floated to the top fraction - consistent with lipid droplet association. However, under the conditions we tested only a small fraction of total protein associated with lipid droplets in either case. MSMEG_2329 may partition between the cytoplasm and lipid droplets while CmrA may associate with both lipid droplets and the PM-CW. It remains to be determined why fluorescence imaging of these proteins revealed the patterns that are similar to those of IMD-associated proteins. It should also be noted that these fractionation patterns of the fusion proteins could be artifacts of fusion constructs rather than representing the true subcellular localization of these proteins. We must await future studies to test these hypotheses for each of these proteins.

How many IMD-associated proteins could there be in *M. smegmatis* genome? Considering that we found three new proteins from screening 523 proteins, we predict that an additional ~30 proteins might be IMD associated (based on a genome encoding 6716 proteins). However, we must also consider the possibility that the list of proteins in the IMD proteome contains additional false positives. Thus, the total number of IMD-associated proteins is likely to remain around 300 proteins. This represents 4-5% of total proteins in *M. smegmatis*. In a broader perspective, 2%, 6%, and 11% of proteins in human cells associate with the endoplasmic reticulum, Golgi apparatus and plasma membrane, respectively (31). Therefore, the estimated percentage of IMD-associated proteins in *M. smegmatis* is reasonable for proteins associating with a subcellular membrane compartment.

In conclusion, the combination of biochemical and microscopic approaches described here has increased the rigor of assigning proteins to a subcellular factory, the mycobacterial IMD. The Mycobacterial Systems Resource offers a valuable platform to integrate knowledge on the atlas of proteins in mycobacterial cells.

## Materials and Methods

### Construction of expression vectors and creation of fluorescence image library

Expression vectors were constructed as a part of the Mycobacterial Systems Resource project (http://msrdb.org/), to be published elsewhere, creating plasmids to express a Dendra2-FLAG fusion of 1118 *M. smegmatis* proteins that represent the core genome conserved across five species of mycobacteria. We analyzed fluorescence images of 523 strains that produced sufficient quality images in the preliminary stage of image acquisition (**Table S1**). Fluorescence localization was categorized as “IMD-like” by satisfying both of the following two criteria: 1) intense enrichment in the polar regions of the cell and 2) less intense sidewall patches. Images of 523 strains were visually screened for IMD-like fluorescent localization.

### Preparation of crude cell lysates

Cells were grown in Middlebrook 7H9 broth supplemented with 11 mM glucose, 14.5 mM NaCl, 0.05% Tween-80, and 12.5 μg/ml apramycin at 30°C with shaking until reaching a log phase (OD_600_ = 0.5-1.0). Cells were spun at 3220x *g* at 4°C, washed in 50 mM Hepes/NaOH (pH 7.4) and resuspended in a lysis buffer containing 25 mM Hepes/NaOH (pH 7.4), 15% glycerol, 2 mM EGTA and a protease inhibitor cocktail (ThermoFisher). Cells were lysed by bead-beating (Millipore-Sigma, acid-washed, 106 μm) and lysate was stored frozen at −20°C.

### BCA Assay

Protein concentration was determined following the manufacturer’s instruction (ThermoFisher) using bovine serum albumin as a standard. Each sample was analyzed in duplicate and the average was used to calculate the protein concentration.

### SDS-PAGE and western blot

Cell lysates were denatured in a reducing sample loading buffer containing 62 mM Tris-HCl (pH 6.8), 50 mM DTT, 2% SDS (w/v) 12.5% glycerol (w/v), and 100 μg/ml Bromophenol Blue at 95°C for 5 min and analyzed by SDS-PAGE (10% gel) using a running buffer containing 25 mM Tris-HCl (pH 8.3), 192 mM glycine, and 0.1% SDS (w/v). Precision Plus Protein Kaleidoscope (BioRad) was used as a molecular weight standard. Proteins were transferred to an Immun-Blot PVDF membrane (BioRad), blocked in 5% skim milk in phosphate-buffered saline (PBS) supplemented with 0.05% Tween-20 (PBST20), and incubated with primary antibodies (mouse anti-FLAG, Millipore-Sigma; rabbit anti-PimB’ (32); rabbit anti-MptA (32); or rabbit anti-MptC (32)). Membrane was washed with PBST20, then incubated with secondary antibodies (horseradish peroxidase-conjugated anti-mouse IgG, GE Healthcare; or horseradish peroxidase-conjugated anti-rabbit IgG, GE Healthcare). Membrane was washed with PBST20 and imaged by chemiluminescence using ImageQuant LAS-4000 mini (GE Healthcare). For sucrose density gradient fractions, an equal volume of each fraction was loaded into each well. Western blot bands were quantified by ImageQuant TL (GE Healthcare) and categorized as follows: Fr. 1-2, cytoplasm; Fr. 3-6, IMD; Fr. 8-11; and Fr. 7/12, other.

### Semi-native PAGE and fluorescence

Protein samples were incubated with non-reducing SDS-free sample loading buffer, containing 62 mM Tris-HCl (pH 6.8), 12.5% glycerol (w/v), and 100 μg/ml Bromophenol Blue, on ice for 30 minutes. The non-denatured samples were analyzed by standard SDS-PAGE (10% gel) in a running buffer containing 25 mM Tris-HCl (pH 8.3), 192 mM glycine, and 0.1% SDS (w/v). Green fluorescence of Dendra2 was visualized using ImageQuant LAS-4000 mini (GE).

### Sucrose density gradient fractionation

Sucrose density gradient fractionation was done as described before (13). Briefly, log phase cells were washed in 50 mM Hepes/NaOH (pH 7.4) and resuspended in a lysis buffer containing 25 mM Hepes (pH 7.4), 20% sucrose, 2 mM EGTA and a protease inhibitor mix (ThermoFisher). Nitrogen cavitation was repeated three times with 1850 - 2200 psi for 30 minutes each. For sedimentation, lysates were placed on top of a 20-50% sucrose gradient and sedimented at 35,000 rpm (218,000x *g*) for 6 hours at 4°C on SW40Ti rotor (Beckman). For flotation, 600 mg of sucrose was dissolved in 900 μl of lysate on a rotating wheel for 2 hours. Lysate was then placed at the bottom of a 20-50% sucrose gradient, and centrifuged at 35,000 rpm (218,000x g) for 18 hours at 4°C. One ml fractions were collected using a gradient collector (BioComp) and stored at - 80°C.

### Electroporation of plasmid constructs

An *M. smegmatis* strain expressing HA-mCherry-GlfT2 from the endogenous locus (13) was electroporated with plasmid DNA at 2.8 kV, 99 μsec, 5 times at 1 sec intervals using a square wave electroporation system (BTX ECM830).

### Fluorescence microscopy and image analysis

*M. smegmatis* cells co-expressing HA-mCherry-GlfT2 and CcdA-Dendra2-FLAG were grown in Middlebrook 7H9 broth to log phase (OD_600_ = 0.5-1.0), and 10 μl of cells were placed onto a 1% agar pad made of Middlebrook 7H9 broth. All images were taken using a fluorescence microscope (Nikon Eclipse E600) with a 100X Plan Fluor 1.30 (oil) objective lens, equipped with a cooled CCD Spot-RT digital camera (Diagnostic Instruments). Cell contours were outlined and fluorescence intensities along the long axis of the cell were measured using Oufti and a custom-written MATLAB codes as described (12, 33). The cell length was normalized by taking the length of the long axis of the cell as 100%. Cells were oriented so that the brighter pole is on the right-hand of the graph.

## Acknowledgements

This work was supported by NIH R03 AI140259 to YSM and NIH R01 AI097191 to KMD, SMF, TAG, and JTW. CAZR was a recipient of the American Society for Microbiology Undergraduate Research Fellowship. JP was a recipient of the Science Without Borders Fellowship from CAPES-Brazil (0328-13-8).

